# Potent protease inhibitors of deadly lagoviruses: rabbit hemorrhagic disease virus and European brown hare syndrome virus

**DOI:** 10.1101/2022.01.10.474982

**Authors:** Krishani Dinali Perera, David Johnson, Scott Lovell, William Groutas, Kyeong-Ok Chang, Yunjeong Kim

**Author notes:** Corresponding author ^1^Department of Diagnostic Medicine & Pathobiology, College of Veterinary Medicine, Kansas State University, Manhattan, KS 66506.

## Abstract

Rabbit hemorrhagic disease (RHD) and European brown hare syndrome (EBHS) are highly contagious diseases caused by lagoviruses in the *Caliciviridae* family and mainly affect rabbits and hares, respectively. These infectious diseases are associated with high mortality and a serious threat to domesticated and wild rabbits and hares, including endangered species such as Riparian brush rabbits. In the US, only isolated cases of RHD had been reported until Spring 2020. However, RHD caused by RHD type 2 virus (RHDV2) was unexpectedly reported in April 2020 in New Mexico and has subsequently spread to several US states infecting wild rabbits and hares, making it highly likely that RHD will become endemic in the US. Vaccines are available for RHD, however, there is no specific treatment for these diseases. RHDV and EBHSV encode a 3C-like protease (3CLpro), which is essential for virus replication and a promising target for antiviral drug development. We have previously generated focused small molecule libraries of 3CLpro inhibitors and demonstrated the *in vitro* potency and *in vivo* efficacy of some protease inhibitors against viruses that encode 3CLpro including caliciviruses and coronaviruses. Here we established the enzyme and cell-based assays for these uncultivable viruses to determine the *in vitro* activity of 3CLpro inhibitors, including GC376, a protease inhibitor being developed for feline infectious peritonitis, and identified potent inhibitors of RHDV1 and 2 and EBHSV. In addition, structure-activity relationship study and homology modelling of the 3CLpros and inhibitors revealed that lagoviruses share similar structural requirements for 3CLpro inhibition with other caliciviruses.

## 1. Introduction

Rabbit hemorrhagic disease (RHD) is a highly contagious viral disease of domesticated (farmed and pet) and wild rabbits with high mortality, which can reach up to 70~100% (Gleeson and Petritz, 2020). RHD is caused by rabbit hemorrhagic disease virus (RHDV) type 1 or 2, which belong to the genus *Lagovirus* in the *Caliciviridae* family (Fig. 1A) (Adams et al., 2017). The classical type 1 RHDV (RHDV1) was first reported in 1984 in China among rabbits imported from Germany, and its variant type 2 RHDV (RHDV2) emerged in 2010 in France (Le Gall-Recule et al., 2011). Since then, RHDV type 1 and 2 infections have become endemic in most parts of the world, including Australia, New Zealand, parts of Asia and most of Europe (Cancellotti and Renzi, 1991), and in some regions RHDV2 mostly replaced RHDV1. In the US, only sporadic, isolated incidences of RHD had been reported in domesticated rabbits until recently (J.A.V.M,, 2000). However, RHDV2 infections in wild black-tailed jackrabbit and wild cottontails were confirmed in New Mexico in April 2020 (New Mexico Department of Game and Fish, 2020) and have been spreading to multiple states among domesticated and wild rabbits, significantly diminishing the hope of eradication of RHD in the US. The characteristic clinical signs of RHD caused by RHDV1 and 2 are fulminant viral hepatitis; The affected animals exhibit lethargy, inappetence and fever followed by death or die before showing clinical signs. Although both RHDV types have similar pathogenesis and clinical course, RHDV2 can infect young (< 2-month-old) and adult rabbits and several hare species (Buehler et al., 2020; Dalton et al., 2012; Le Gall-Recule et al., 2017; Velarde et al., 2017), while RHDV1 infection is limited to adult domestic rabbit populations. RHDV also pose a serious threat to threatened or endangered wild rabbit species, such as Riparian Brush rabbits, Riverine rabbit, Pygmy rabbits or Hispid hare (also called as bristly rabbits) (Villafuerte and Delibes-Mateos).

**Figure 1.**
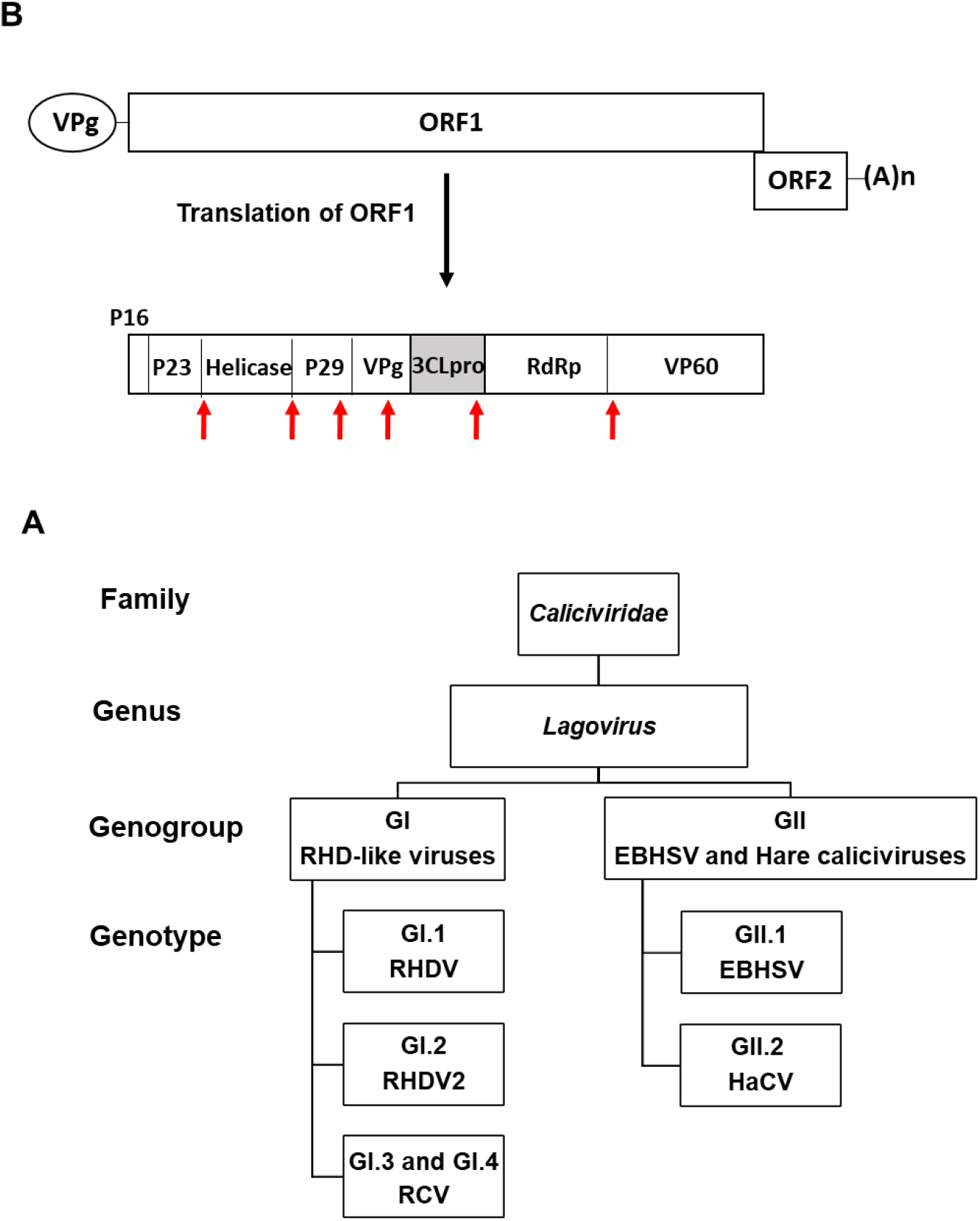
Classification and genomic organization of RHDV and EBHSV. **(A)** The *Caliciviridae* family contains 11 genera including *Lagovirus* genus. Two main genogroups GI and GII in the *Lagovirus* genus further divide into several genotypes. GII.2 is the putative genotype that contains hare caliciviruses (Le Pendu *et al,* 2017). **(B)** Genome organization of RHDV and EBHSV. The genome is composed of two overlapping ORF1 and ORF2. ORF1 translates into a polyprotein which is proteolytically cleaved to generate nonstructural proteins and the capsid (VP60) and ORF2 codes for a minor structural protein VP10. The red arrows indicate the cleavage site on the virus polyproteins that are processed by 3CLpro. The genomic and subgenomic RNA (not shown) have a virus-encoded protein VPg attached to their 5’end and are polyadenylated at their 3’end. RHD, rabbit hemorrhagic virus; RCV, rabbit calicivirus; EBSHV, European brown hare syndrome virus; HaCV, hare calicivirus. ORF, open reading frame; RdRp, RNA-dependent RNA polymerase.

European brown hare syndrome virus (EBHSV), another lagovirus (Fig. 1A), is a highly pathogenic and contagious virus that mainly affects hares, such as European brown hares, mountain hares and Italian hares, although the eastern cottontail was reported to be susceptible to spillover infection of EBHSV (Velarde et al., 2017). EBSHV was first detected in Sweden in the early 1980s and has since become endemic in Europe (Gavier-Widen and Morner, 1993). EBSHV causes fatal viral hepatitis with clinical signs similar to those of RHDV, and the mortality rate of EBSH may reach 100% in farmed hares (Abrantes et al., 2012). Rabbit calicivirus (RCV) and hare calicivirus (HaCV) are non-pathogenic members of the genus *Lagovirus* (Fig. 1A) and have been shown to undergo recombination with RHDV and EBHSV contributing to strain diversity and evolution of viruses (Abrantes et al., 2020; Szillat et al., 2020).

These caliciviruses are highly contagious, requiring a small number of virus particles for infection, and very stable in the environment. It was reported that RHDV can survive for up to three months in rabbit carcass (Henning et al., 2005), and bodily fluid, fomites and biting insects are able to spread the virus as mechanical vectors (McColl et al., 2002). Furthermore, surviving rabbits can shed RDHV for months. Therefore, it is very difficult to control viral transmission once it is spread to wild rabbit populations. Although no vaccine is available for EBSHV, inactivated and myxoma virus-based live vaccines are licensed for RHDV in the EU. In the US, recombinant subunit vaccine for RHDV2 was recently issued emergency use authorization by USDA’s Center for Veterinary Biologics in October 2021. However, no specific treatment is available for RHDV and EBSHV, and only strict biosecurity practices and vaccinations are used for prevention and control of the viruses. Considering the detrimental impact of these virus infections on the global rabbit and hare industry and pet rabbits as well as endangered rabbit and hare species, development of effective prophylactic and therapeutic treatment would be a valuable contribution to the limited arsenal of control measures available against these virus diseases.

Caliciviruses such as RHDV and EBSHV have a single-stranded, positive sense RNA genome, which is organized into two major open reading frames (ORFs) (Fig. 1B). The ORF1 encodes a polyprotein, which is processed by virus-encoded 3C-like protease (3CLpro) to release mature nonstructural proteins and the major structural protein VP1. This virus polyprotein processing by 3CLpro is an essential step in virus replication and is thus considered a promising target for drug discovery for viruses that encode 3CLpro, such as caliciviruses and coronaviruses. We have previously reported potent 3CLpro inhibitors of caliciviruses including human norovirus (Damalanka et al., 2016; Damalanka et al., 2017; Damalanka et al., 2018; Galasiti Kankanamalage et al., 2017a; Galasiti Kankanamalage et al., 2015; Galasiti Kankanamalage et al., 2019; Kim et al., 2012; Mandadapu et al., 2013a; Mandadapu et al., 2013b; Mandadapu et al., 2012; Rathnayake et al., 2020a; Weerawarna et al., 2016) and feline calicivirus (Kim et al., 2015) and of coronaviruses such as feline infectious peritonitis virus (FIPV) (Kim et al., 2016; Pedersen et al., 2018), MERS-CoV (Rathnayake et al., 2020b), MERS-CoV (Rathnayake et al., 2020b) and SARS-CoV-2 (Dampalla et al., 2021a; Dampalla et al., 2021b). In this study, we established a fluorescence resonance energy transfer (FRET) assay and a cell-based reporter assay to evaluate 3CLpro inhibitors and identified potent compounds against RHDV1 and 2 and EBSHV. Structure-activity relationship study and 3D homology modelling of 3CLpros of RHDV1 and 2 and EBSHV was also conducted to understand the structural basis for the potency of the compounds, which showed that structural requirements of compounds for the inhibition of 3CLpros are similar among caliciviruses including lagoviruses.

## 2. Materials and methods

### 2.1. Compounds

The 3CLpro inhibitors including NPI52 (Prior et al., 2013), GC376 (Kim et al., 2012), GC543 (Mandadapu et al., 2013b), GC583 (Galasiti Kankanamalage et al., 2015), GC571 (Galasiti Kankanamalage et al., 2015) and GC772 (Galasiti Kankanamalage et al., 2017b) have been previously generated and reported by us. Table 1 contains the list of the compounds used in the study.

**Table 1.** Structures of the inhibitors used in the assays. The structure of NPI52, a tripeptidyl compound, is also shown.

| Inhibitor |  | Structural moiety |  |  |  |
| --- | --- | --- | --- | --- | --- |
|  |  | R<sub>1</sub> | R<sub>2</sub> | R<sub>3</sub> | R<sub>4</sub> |
| Dipeptidyl | GC376 | (C<sub>6</sub>H<sub>5</sub>)CH<sub>2</sub> | O(C=O)NH | Leu<br>(Isobutyl) | CH(OH)SO<sub>3</sub>N<sub>a</sub> |
|  | GC543 | (C<sub>6</sub>H<sub>5</sub>)CH<sub>2</sub> | O(C=O)NH | Cha<br>(Cyclohexyl<br>methyl) | CHO |
|  | GC583 | m-Cl(C<sub>6</sub>H<sub>4</sub>)CH<sub>2</sub> | O(C=O)NH | Cha | CHO |
|  | GC591 | m-Cl(C<sub>6</sub>H<sub>4</sub>)CH<sub>2</sub> | O(C=O)NH | Cha | (C=O)CONH<br>cyclopropyl |
|  | GC772 | m-Cl(C<sub>6</sub>H<sub>4</sub>) | SO<sub>2</sub>NH | Cha | CHO |

### 2.2. Expression and purification of recombinant RHDV1 and 2 and EBHSV 3CLpros for FRET assay

The 3CLpro sequences of RHDV1 (FRG strain), RHDV2 (NL2016 strain) and EBHSV (G104 strain) were obtained from GenBank (accession numbers NP_062875.1, QDF46325.1 and QED89978.1, respectively). The codon-optimized, full length 3CLpro coding-sequences with N-terminal 6His tags were synthesized by Integrated DNA Technologies (Coralville, IA) and cloned into pET-28a(+) vector (Addgene, Cambridge, MA). Subsequently, the recombinant 3CLpros were expressed in *Escherichia coli* BL21 cells (Invitrogen, Carlsbad, CA) grown in Luria Bertani broth induced with 1mM isopropyl ß-D-thiogalactopyranoside. The recombinant 3CLpro were then purified using HIS Gravitrap Ni-NTA affinity columns (GE Healthcare, Chicago, IL), following the standard protocol (Kim et al., 2012).

### 2.3. The FRET assays

First, activities of the recombinant 3CLpros of RHDV1, RHDV2 and EBHSV were evaluated. Briefly, serial dilutions of 3CLpro were made in 25 μl of assay buffer (50mM NaCl, 6mM Dithiothreitol, 50 mM HEPES, 0.4mM EDTA and 60% Glycerol at pH 8.0) and mixed with the same volume of assay buffer containing the FRET substrate 5-FAM-ASFEGS-K(QXL520)-NH2 (AnaSpec, Fremont, CA). The cleavage site (ASFEGS) of the FRET substrate was derived from the cleavage site of RHDV NSP3/4. The mixture was then added into a black 96 well imaging microplate (Fisher Scientific, Waltham, MA). Following incubation of the plate at room temperature (RT) for up to 90 min, fluorescence readings were measured at excitation and emission values 485 and 516 nm, respectively, on a fluorescence microplate reader (FLx800, Biotek, Winnooski, VT). The relative fluorescence (activity) of each 3CLpro over time was determined by subtracting the readings of substrate only control from raw fluorescence readings. After confirming the activity of 3CLpros, the inhibitory activity of the compounds was determined against 3CLpros as previously described (Kim et al., 2012; Perera et al., 2018). Serial dilutions of each compound (10 mM stock solution in DMSO) were prepared in DMSO or media, and each dilution was added to 25 μl of assay buffer containing the 3CLpro. The concentration of DMSO in each dilution did not exceed 4 % of final concentration. After incubation at RT for 30 min, the mixture was added into 25 μl of assay buffer containing the FRET substrate in a black 96 well microplate, and the plate was incubated at RT for 30 min. The raw fluorescence readings were measured on the fluorescence microplate reader, and the relative fluorescence was calculated by subtracting the values for substrate-only control from the raw values. The 50% inhibitory concentration (IC_50_) was calculated using the non-linear regression analysis (four parameter variable slope) in GraphPad Prism software version 6.07 (GraphPad Software, La Jolla, CA).

### 2.4. Cell-based reporter assays. *Generation of plasmids encoding RHDV2 3CLpro and circular permutated firefly genes*

The full-length 3CLpro sequence of RHDV2 was codon-optimized for protein expression in mammalian cells, synthesized by Integrated DNA Technologies (Coralville, IA) and inserted into pcDNA3 H2B-mIFP T2A Mpro (3CLpro) (Addgene, MA) following digesting with BamH1 and ApaI. The resulting plasmid was designated as pcDNA3-RHDV2-3CLpro. A plasmid encoding inactive RHDV2 3CLpro was generated by substituting the nucleophilic Cys with Ala in the active site of 3CLpro in pcDNA3-RHDV2 using site-directed mutagenesis (Agilent’s Quik Change mutagenesis kit, Agilent, CA) and designated as pcDNA3-RHDV2-m3CLpro. The pGloSensor^TM^-30F-DEVDG plasmid (Promega, Madison, WI) contains firefly luciferase gene with a caspase 3/7 cleavage site (DEVDG), which is a circular, permutated form of firefly luciferase (Galban et al., 2013). To replace the caspase 3/7 cleavage site with a sequence from RHDV NS3/4 cleavage site (VASFEGAN), site-directed mutagenesis was conducted using primers containing the cleavage sequence. The primer sequences are Forward: 5’-GATCCGTGGCCTCATTCGAGGGTGCA-3’ and Reverse: 5’-AGCTTGCACCCTCGAATGAGGCCACG-3’. The resulting plasmid was designated as pGlo-RHDV. The plasmid also contains Renilla luciferase as an expression control. ***Assay***. One-day old 293T cells in a 48-well plate were co-transfected with pGlo-RHDV and pcDNA3-RHDV2-3CLpro or pcDNA3-RHDV2-m3CLpro. About 16 hr later, medium was replaced with new medium containing serial dilutions of each compound or mock (medium), and the cells were incubated for 37°C for 5 hours. Then, the cells were lysed and firefly and Renilla luminescence were measured following the manufacturer’s directions (Dual luciferase kit, Promega, Madison, WI) on a luminometer (GloMax^®^ 20/20 Luminometer, Promega, Madison, WI). Firefly luciferase was normalized against Renilla luciferase, and the 50% effective concentration (EC_50_) of each compound was calculated by GraphPad Prism software using a variable slope (GraphPad, La Jolla, CA). Figure 2 illustrates the cell-based reporter assay.

**Figure 2.**
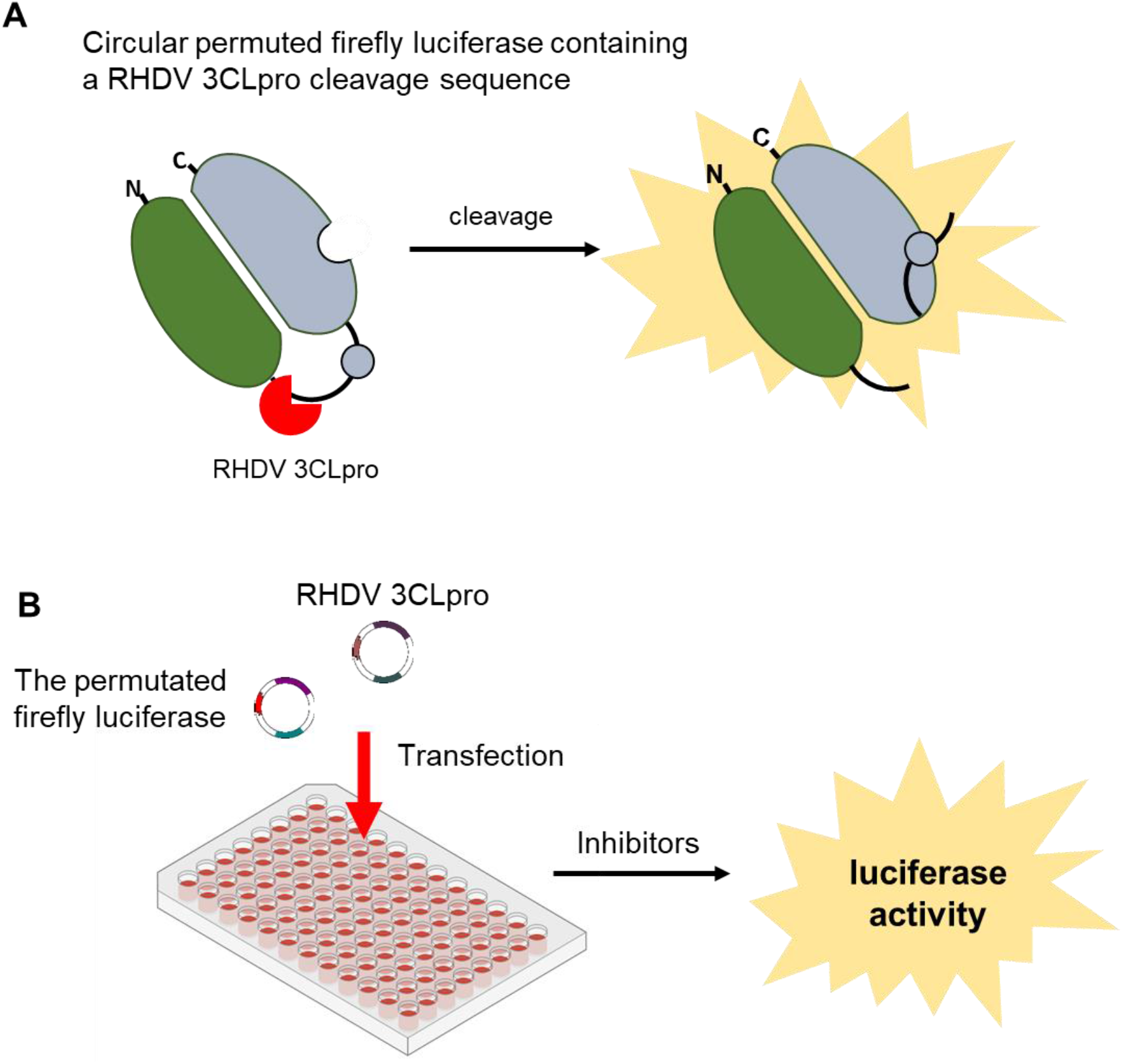
Cell-based reporter assay. **(A)** A circularly permuted form of firefly luciferase containing the RHDV2 3CLpro cleavage sequence was generated by replacing the caspase 3/7 recognition sequence in the pGloSensor-30F plasmid construct. Cleavage of the 3CLpro recognition sequence leads to activation of firefly, resulting in an increase in luminescence activity. **(B)** For inhibition assay, one-day old 293T cells were transfected with plasmids encoding the permuted firefly luciferase and RHDV2 3CLpro genes. After 16 hrs of incubation of the cells, different concentrations of compounds were added to the cells. Then the cells were further incubated for 5 hrs before lysis of the cells. Firefly and Renilla luciferase activities were measured for the determination of the EC_50_ of each compound.

### 2.5. Cytotoxicity assay

To determine the cytotoxicity of the compounds, semi-confluent 293T cells in a 96-well plate were incubated with serial dilutions of each compound (up to 100 μM) for 5 hrs. Cell cytotoxicity was determined by a CytoTox 96 nonradioactive assay kit (Promega, Madison, WI), and the 50% cytotoxic concentration (CC_50_) value for each compound was calculated using GraphPad Prism software. The non-specific cytotoxic effects of these compounds were also reported previously (Galasiti Kankanamalage et al., 2017b; Galasiti Kankanamalage et al., 2015; Kim et al., 2012; Mandadapu et al., 2013b; Prior et al., 2013).

### 2.6. Multiple amino acid sequence alignment and generation of phylogenetic tree of RHDV, EBHSV, RCV and HaCV 3Cpros

The 3CLpro amino acid sequences of RHDV1 (FRG strain), RHDV2 (NL2016 strain), EBHSV (G104 strain), RCV (Gudg-79 strain) and HaCV (E15-431 strain) were obtained from Genbank and aligned using Clustal Omega (https://www.ebi.ac.uk/Tools/msa/clustalo/) (Madeira et al., 2019). To generate a phylogenetic tree, the 3CLpro amino acid sequences of RHDV1 (46 sequences), RHDV2 (39 sequences), EBHSV (6 sequences), RCV (39 sequences) and HaCV (5 sequences) were obtained from Genbank and aligned using Clustal Omega and a phylogenetic tree was generated using the maximum Likelihood method and Whelan and Goldman model (Whelan and Goldman, 2001) in MEGA X (Kumar et al., 2018) and annotated using iTOL v6.4 (Letunic and Bork, 2021).

### 2.7. Three-dimensional structural homology modeling of 3Cpro of RHDV1, RHDV2 and EBHSV

The three-dimensional homology structures of 3CLpro of RHDV1 (FRG strain), RHDV2 (NL2016 strain), and EBHSV (G104 strain) were generated using the Phyre2 web portal (http://www.sbg.bio.ic.ac.uk/phyre2) (Kelley et al., 2015). RHDV1 and RHDV2 3CLpros were modeled using NS6 protease of murine norovirus 1 (Protein Data Bank ID:4ASH) as a template. The templates for modeling EBSHV 3CLpro were a hydrolase of *E. coli* (PDB ID 2ZLE) and a lyase of *Campylobacter jejuni* (PDB ID 6Z05). In general, all the homology models were generated at >90% confidence for 91-100% residues, and TM scores and RMSD values ranged from 0.31-0.62 and 1.86-2.56 Å, respectively (https://zhanggroup.org/TM-align/) (Zhang and Skolnick, 2005). The constructed RHDV1, RHDV2 and EBHSV 3CLpro models were superposed with human norovirus (Genus *Norovirus* in the *Caliciviridae* family) 3CLpro-GC376 co-crystal structure (PDB accession number: 3UR9) using the PyMol molecular graphics system, Version 1.8 (Schrodinger LLC, Cambridge, MA) (DeLano, 2010) or SuperPose Version 1.0 (http://superpose.wishartlab.com/) (Rajarshi Maiti, 2004) to compare the active site conformations. Inhibitor bound models of RHDV2 3CLpro with GC376 and GC583 were generated using Alphafold2 (Jumper et al., 2021) and by aligning the binding sites of structures, with the highest sequence similarity that included human norovirus 3CLpro bound with GC376 [PDB 3UR9 (Kim et al., 2012)] and GC583 [PDB 4XBB (Galasiti Kankanamalage et al., 2015)]. The ligand bound structure prediction was optimized by superimposing the ligand from the aforementioned crystal structures, adding the covalent bond with the active site cysteine and running Schrodinger’s protein preparation wizard to optimize hydrogen bonding and minimize the structure into Schrodinger’s OSPL4 energy function (Schrödinger LLC, 2019).

## 3. Results

### 3.1. Inhibitory activity of compounds against RHDV1 and 2 and EBHSV 3CLpros in the FRET assay

Activities of the recombinant RHDV1 and 2 and EBHSV 3CLpro were confirmed prior to inhibition assay. Each 3CLpro showed gradual increase in activity over time in a similar trend (Fig. 3A). Next, we evaluated the compounds against each 3CLpro in the assay to determine the IC_50_ values against each 3CLpro (Fig. 3B). These compounds have the same skeletal structure with a glutamine surrogate at the P1 position and various functional groups at R1-R3 and different reactive warheads (R4) that interact with the cysteine residue at the catalytic site. Among the tested compounds, GC376 showed a lower inhibitory activity against all 3CLpros compared to other tested compounds with 1.21, 1.39 and 1 μM IC_50_ values against RHDV1, RHDV2 and EBHSV 3CLpro. The IC_50_ value of GC376 against feline coronavirus 3CLpro has been previously reported to be about 0.5 μM in the FRET assay by us (Perera et al., 2018). GC376 is a dipeptidyl compound that potently inhibits various coronaviruses (Kim et al., 2016; Perera et al., 2018) and contains Leu and benzyl ring at the R_3_ and R_1_ positions, respectively, and a bisulfite adduct warhead (R_4_) that converts to an aldehyde form (Table 1) (Kim et al., 2016). Substitution of the bisulfite adduct warhead and Leu at the R3 (GC376) with an aldehyde warhead and Cha (GC543) led to greatly increased inhibitory activity against all 3CLpros. We have previously showed that compounds with bisulfite adduct vs aldehyde warheads exhibit similar activity in the FRET and cell-based assays (Galasiti Kankanamalage et al., 2017b). Therefore, substitution of Leu with Cha at the R_3_ would be responsible for the observed increase in potency. Replacing benzyl ring (GC543) with m-benzyl ring (GC583) at R_1_ slightly further increased potency against all tested 3CLpros. However, replacing the aldehyde warhead (GC583) with (C=O)CONH cyclopropyl (GC591) markedly decreased the inhibitory activities against all 3CLpros, but this reduction in activity was not as profound as having Leu at R_3_ (GC376). In GC772, the carbamate moiety of GC583 was replaced with a sulfonamide linkage, which decreased potency against all tested 3CLpros. NPI52, a tripeptidyl compound, has homologous structural elements with GC376 except that NPI52 has an aldehyde warhead and an additional residue of 1-naththylalanine that corresponds to the P3 position. The inhibitory activity of NPI52 was more potent than GC376 but less than GC543 and GC583 against the tested 3CLpros. Dose-response curves of GC589 against RHDV1 and 2 and EBHSV 3CLpros are shown in Fig. 3C.

**Figure 3.**
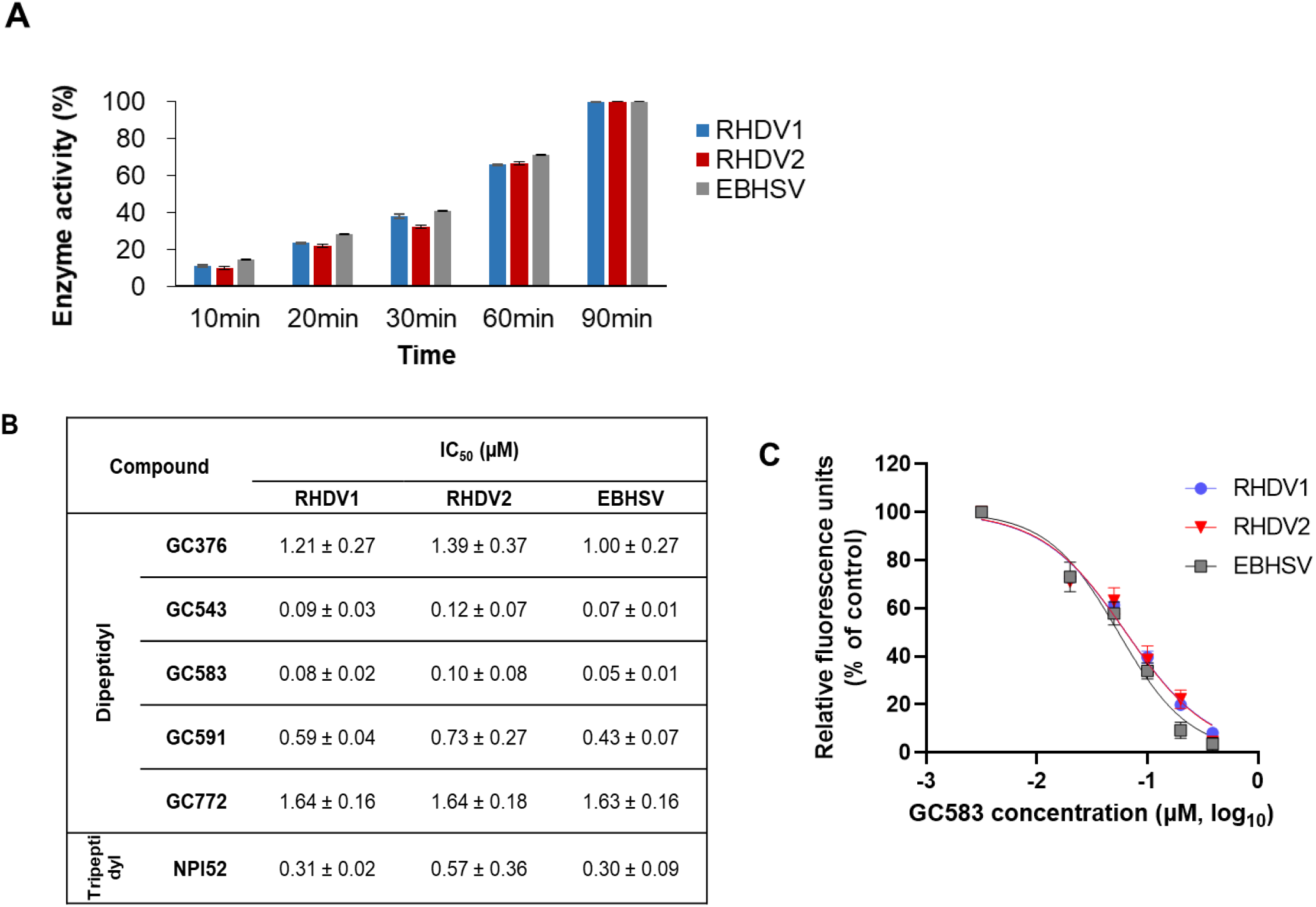
Activity of the compounds against 3CLpros of RHDV1, RHDV2 and EBHSV in FRET assay. **(A)** Activities of the recombinant RHDV1, RHDV2 and EBSHV 3CLpros. Briefly, the recombinant 3CLpro was added to the assay buffer containing the fluorescence substrate and the mixture was incubated at room temperature. The fluorescence measurements were measured for up to 90 min and the percentage activity of each 3CLpro was calculated. **(B)** The activity of compounds against RHDV1, RHDV2 and EBSHV 3CLpros in the FRET assay. Briefly, serial dilutions of each compound were added to assay buffer containing 3CLpro. After 30 min of incubation, the mixture was added to assay buffer containing the FRET substrate. Raw fluorescence readings were obtained 30 min later, and relative fluorescence was calculated by subtracting substrate-only control from raw values to calculate the 50% inhibitory concentration (IC_50_) for each compound. **(C)** Dose dependent inhibition curves of GC583 against RHDV1, RHDV2 and EBSHV 3CLpros.

### 3.2. Inhibitory activity of the compounds in a cell–based luciferase reporter assay

Since lagoviruses do not grow in cell culture, we examined the potency of the compounds against RHDV2 in a cell-based reporter assay (Fig. 2A and B). The inactive RHDV 3CLpro was also included as a control and had minimal luminescence in the assay (<1% of luminescence of active 3CLpro). All the compounds that were included in the study showed minimal cytotoxicity at 100 μM in 293T cells (Fig. 4A). The EC_50_ values of these compounds against RHDV2 3CLpro ranged from 0.46 ~ 9.80 μM in 293T cells (Fig. 4A). In this assay system, GC543 and GC583 exhibited most potent activity, which is consistent with the results from the FRET assays (Fig. 3B). GC376 and GC772 showed substantially reduced activity compared to GC543 or GC583 by 9.7~11.6-fold, which is comparable to the fold reductions in potency observed in the FRET assay (13.4~15.1-fold). In general, GC591, GC772 and NPI52 showed similar trend of activity compared to GC376 in the cell based and FRET assays except for GC591, which had somewhat decreased activity in the cell-based assay. The dose-dependent curves of GC543 and GC583 against RHDV2 in the cell-based assay are shown in Fig. 4B.

**Figure 4.**
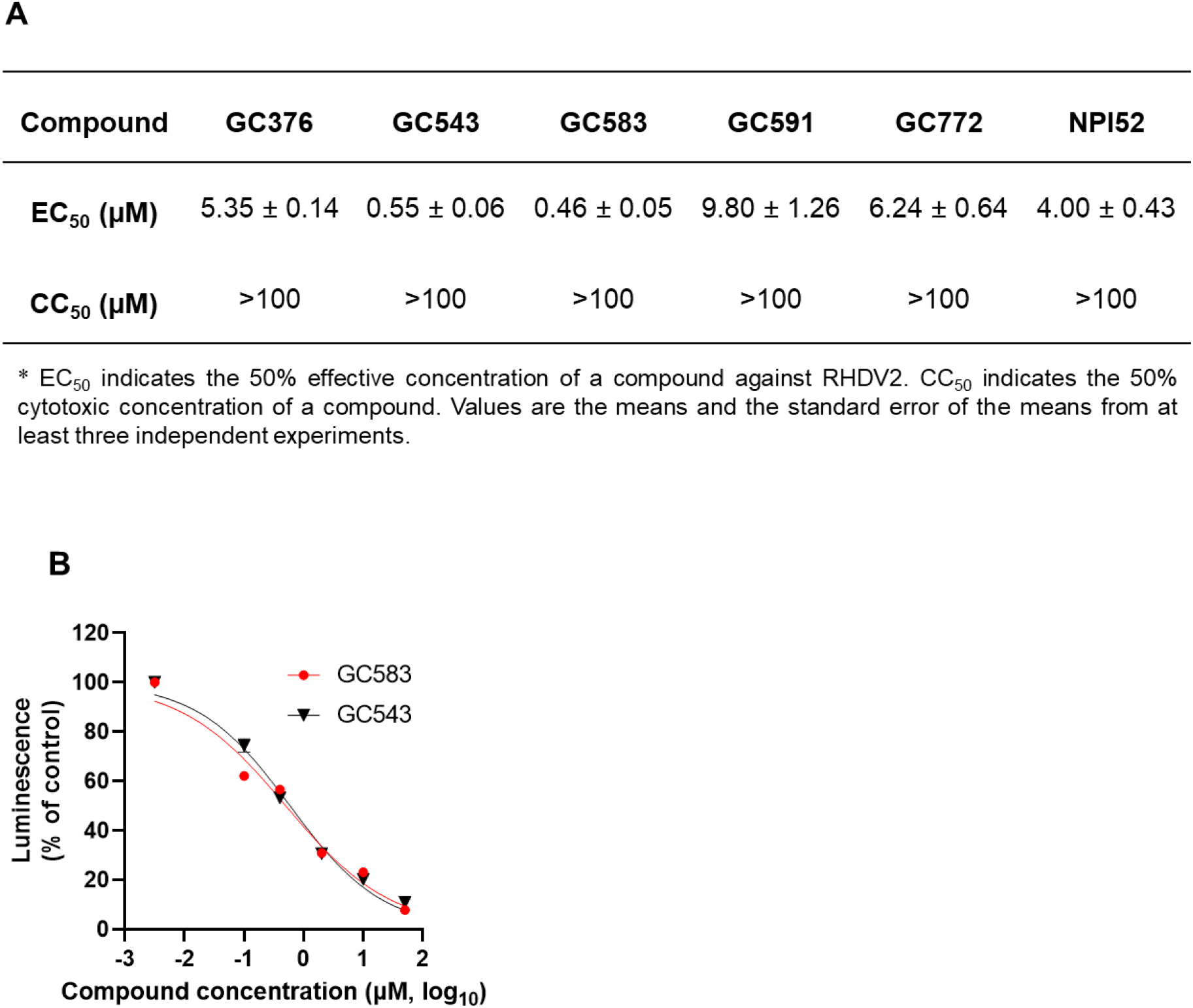
Activity of the compounds against 3CLpros of RHDV1, RHDV2 and EBHSV in cell-based reporter assay. (A) RHDV2 3CLpro inhibitory activity in cell-based luciferase reporter assay and cytotoxicity of the compounds. (B) Dose dependent inhibition curve of GC583 based on percent luciferase activity by RHDV2 3CLpro in 293T cells (R2 > 0.9).

### 3.3. Amino acid sequence homology, multiple amino acid sequence alignment and phylogenetic tree of 3CLpro of RHDV, EBHSV, RCV and HaCV

The amino acid sequence homology of RHDV, EBHSV, RCV and HaCV was investigated. The amino acid sequence homology within each virus was high ranging from 92.31 ~100% for RHDV1, 93.01 ~100% for RHDV2, 97.90~100% for EBHSV, 96.50~100% for RCV, and 86.01~94.41% HaCV. The close genetic relationship of 3CLpros shared among RHDV1, RHDV2 and RCV (GI lagoviruses) was evident with sequence homology of 93.01~100% between RHDV1 and RHDV2, 93.01 ~97.90% between RCV and RHDV1 and 93.01~98.60% between RCV and RHDV2. The 3CLpro sequences of EBSHV and HaCV (GII lagoviruses) were closely related to each other with amino acid sequence homology of 86.01~96.50%. However, as expected, the 3CLpro amino acid sequences between GI and GII lagoviruses are less conserved; EBHSV and HaCV 3CLpros sequences have 82.52~88.11% homology to those of RHDV1 or 2 or RCV. Despite the diversity of 3CLpro sequences among lagoviruses, multiple amino acid sequence alignment of 3CLpros showed that the catalytic residues H27, D44 and C104 (Oka et al., 2011) are conserved in RHDV, EBHSV, RCV and HaCV 3CLpros (Fig. 5A). Phylogenetic analysis showed the 3CLpros of GII lagoviruses, EBHSV and HaCV, can be clustered into the same main branch, while the 3CLpros of GI lagoviruses, RHDV1 and 2 and RCV, belonged to other branches (Fig. 5B). Interestingly, the clades of 3CLpros of GI lagoviruses, RHDV1 and 2 and RCV, are interspersed phylogenetically, which may reflect recombination events that have occurred among these pathogenic and non-pathogenic lagoviruses. It has been reported that lagoviruses have high capacity of recombination within lagoviruses with hotpots between the structural and nonstructural genes (Abrantes et al., 2020; Mahar et al., 2021; Szillat et al., 2020), which is frequently observed for other caliciviruses (Bull et al., 2005; Ludwig-Begall et al., 2018)

**Figure 5.**
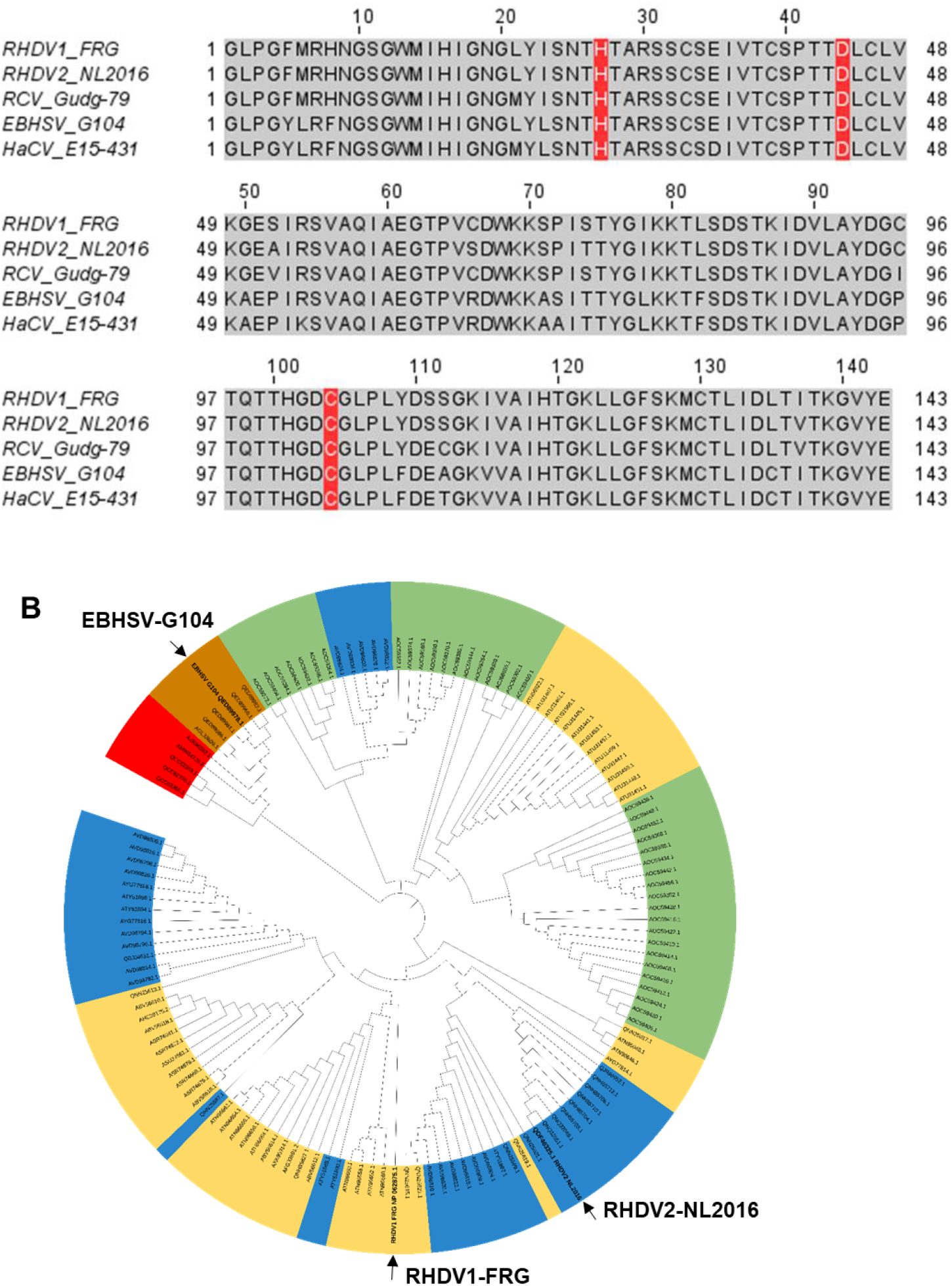
Multiple sequence alignment and phylogenetic analysis of lagovirus 3CLpros. (A) Multiple sequence alignment of 3CLpros of the representative strains of RHDV1, RHDV2, RCV, EBHSV and HaCV. The catalytic residues of 3CLpros (H27, D44 and C104) are highlighted in red. (B) A phylogenetic tree of amino acid sequences of lagovirus 3CLpros, which were retrieved from GenBank was generated using Maximum Likelihood method and Whelan and Goldman model in MEGA X and annotated using iTOL v6.4. Each entry indicates GenBank accession number and is colored yellow (RHDV1), blue (RHDV2), green (RCV), brown (EBHSV) or red (HaCV). The 3CLpros used in the assays are indicated by black arrows.

### 3.4. Three-dimensional homology structural models for 3CLpro of RHDV and EBHSV 3CLpro

The homology-based 3CLpro structural models of RHDV1, RHDV2, and EBHSV are shown in figure 6A and B with the catalytic residues indicated in red. RHDV1 and RHDV2 3CLpro models are very closely homologous (Fig. 6A) with a root-meansquare deviation (RMSD) of 0.24 Å between the Cα atoms of residues 27-104 containing the catalytic residues. However, the 3D model of EBSHV 3CLpro showed a lower structural homology with RHDV1 and RHDV2 3CLpro (Fig. 6B) with a RMSD of 9.9 Å between the Cα atoms from residues 27-104. A crystal structure of human norovirus 3CLpro-GC376 complex was included as its overall structural folds resembles those of lagoviruses (Fig. 6C). The nucleophile residue (C139) of human norovirus forms a covalent bond with GC376 superimposes with the catalytic residues of lagoviruses (Fig. 6D). Despite some differences in the overall structures in these models, the catalytic residues in the catalytic site of all three models align very closely (Fig. 6D). High-confidence models of the 3CLpro of RHDV2, which adopts a similar fold to other structures such as human norovirus 3CLpro (Fig. 6A and C) were docked with GC376 and GC583 to compare the binding modes. The GC376 inhibitor forms hydrogen bond interactions with H27, T99, H101, H119, T120 and K122 (Fig. 7A). Similarly, GC583 forms interactions with H27, T99, H101, H119, and K122 (Fig. 7B) but does not interact with T120. The main difference is observed in the P2 position of the inhibitors where the cyclohexyl group of GC583 occupies the *S2* subsite that is formed by H27, D44 and L82. The benzyl and m-chlorobenzyl rings of GC376 and GC583 respectively are positioned in a hydrophobic cleft in the S4 subsite formed by T81, A92 and L124 (Fig. 7C and D).

**Figure 6.**
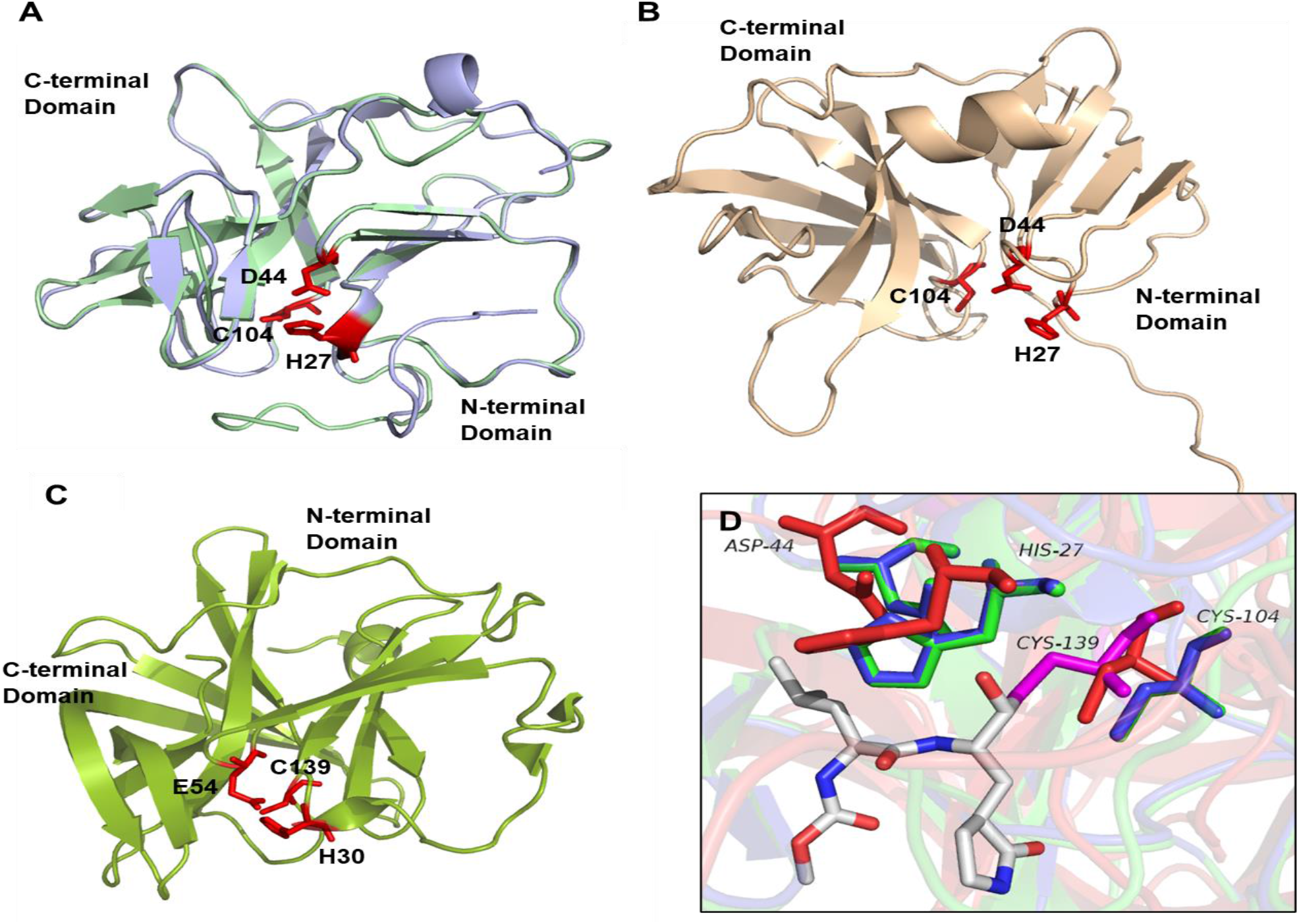
Three-dimensional homology structural models of RHDV1, RHDV2 and EBHSV 3CLpro. **(A)** RHDV1 (pale green) and RHDV2 (pale blue) 3CLpro models are superposed. The catalytic residues (red) overlap each other closely. **(B)** The 3D model of EBHSV 3CLpro shows low structural homology to RHDV1 or RHDV2 3CLpro models. The catalytic residues of EBSHV 3CLpro are also shown in red. **(C)** The crystal structure of human norovirus 3CLpro (Protein Data Bank: 3UR9) with active site residues (red). **(D)** The catalytic site of superposed human norovirus 3CLpro-GC376 complex (Protein Data Bank: 3UR9) and 3D models of RHDV1 (green), RHDV2 (blue) and EBHSV (red) 3CLpro. GC376 is shown as a stick in the catalytic site and forms a covalent bond with the nucleophilic residue (C139) of human norovirus 3CLpro (magenta). The nucleophilic residue (C104) of lagovirus 3CLpros [blue, green (overlapped with blue) or red] are closely aligned. The other catalytic residues of Lagovirus 3CLpro (H27 and D44) are also closely aligned.

**Figure 7.**
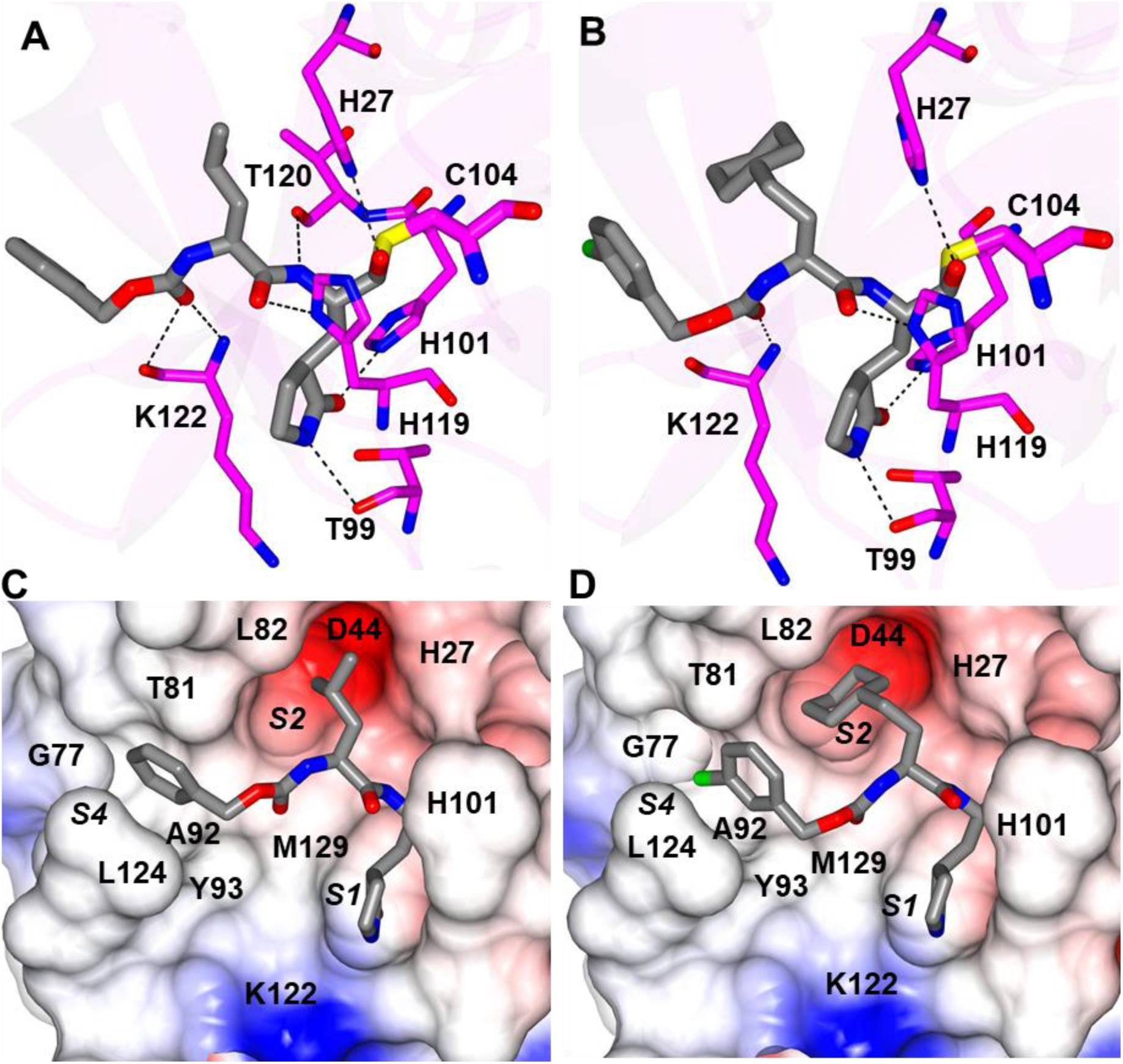
Inhibitors GC376 (**A/C**) and GC583 (**B/D**) docked to RDHV2 3CLpro. Hydrogen bond interactions (dashed lines) are shown in panels A and B and the electrostatic surface representations in panels C and D. The *S1, S2* and *S4* subsites are indicated in parenthesis.

## 4. Discussion

Following the unexpected report of RHDV2 infection in native North American jackrabbits and cottontails in New Mexico in 2020 April, this highly pathogenic calicivirus has been confirmed in wild rabbits and hares in the southwest states in the US (U.S. Food and Drug Administration, Nov 19, 2020) and is now considered endemic or near endemic in these regions. Before this recent spread of RHDV2 in wild rabbits and hares, detection of RHDV2 infection had been limited to isolated cases of domestic and feral rabbits in the US. Outside the US, RHDV1 and subsequently RHDV2 have become endemic in many parts of the world with a substantial economic and ecological impact on the rabbit industry and wild rabbit populations worldwide. However, efforts to investigate potential antiviral treatments have been generally lacking, and specific treatment options are not available for RHD as well as EBHS, which is also endemic in most parts of the world (Du et al., 2019; Urakova et al., 2016). Caliciviruses, coronaviruses and picornaviruses encode viral proteases 3CLpro or 3Cpro, which are responsible for virus polyprotein cleavage and an essential component of virus replication. These viral proteases adopt chymotrypsin-like protease folds and are functionally and structurally conserved among calicivirus, coronavirus and picornaviruses although protein sequence homology may vary substantially among them. We have previously generated focused libraries of 3CLpro/3Cpro inhibitors and reported the *in vitro* and/or *in vivo* potency of some protease inhibitors against caliciviruses, coronaviruses and picornaviruses (Dampalla et al., 2021a; Kim et al., 2016; Kim et al., 2012; Kim et al., 2015; Rathnayake et al., 2020a; Rathnayake et al., 2020b). In this study, we evaluated the activity of select protease inhibitors including GC376 and its derivatives against RHDV1 and 2 and EBHSV 3CLpros and conducted structure-activity relationship studies using the FRET and cell-based reporter assays.

All the compounds we tested showed potent activity against RHDV1 and 2 and EBHSV 3CLpros in the FRET and the cell-based reporter assays, and the activity of each compound was comparable among RHDV1, RHDV2 and EBHSV 3CLpro. GC376 is a dipeptidyl compound that was shown to be broadly effective against various coronaviruses, including FIPV, a highly virulent feline coronavirus, MERS-CoV, SARS-CoV-2, as well as some caliciviruses and picornaviruses (Kim et al., 2016; Kim et al., 2012; Kim et al., 2015; Pedersen et al., 2018; Perera et al., 2018; Vuong et al., 2020) and is currently under commercial development as an antiviral drug for FIP. GC376 is a bisulfite adduct prodrug of the corresponding parent aldehyde GC373 (Kim et al., 2016) and has glutamine surrogate, Leu and benzyl group in the P1, P2 (R_3_ group) and P3 (R_1_ group) position, respectively (Table 1). Bisulfite adduct compounds convert to their aldehyde counterparts, and they exhibit similar potency in *in vitro* assays (Kim et al., 2013; Mandadapu et al., 2013b; Perera et al., 2018). We have previously reported that the antiviral activity of GC376 is slightly more potent against FIPV compared to aldehyde compounds GC543 and GC583, which have Cha (cyclohexylalanine) moiety at R_4_ and a benzyl group or m-chlorobenzyl group at R_1_, respectively (Perera et al., 2018). In contrast, GC543 and GC583 have significantly improved activity against RHDV1 and 2 and EBHSV 3CLpros over GC376, which indicates that the Cha moiety at R_3_ is more suitable for these 3CLpros. The preference of Cha moiety over Leu at R_3_ was also observed for 3CLpro of human norovirus, a calicivirus in the *Norovirus* genus (Galasiti Kankanamalage et al., 2015) and feline calicivirus that belongs to the *Vesivirus* genus (Kim et al., 2015). However, substitution of the benzyl group (GC543) with m-chlorobenzyl (GC583) did not markedly increase the activity of a compound against RHDV1 and 2 and EBHSV 3CLpros, unlike human norovirus 3CLpro. The results from the cell-based reporter assays we have established for RHDV and EBHSV correlated well with those determined in the FRET assay. The EC_50_ values from the reporter assay were generally higher than the IC_50_ values from the FRET assay, which is likely due to transient expression of high levels of 3CLpro in cells. One of the protease inhibitors tested, GC376, has an IC_50_ value of 0.5 μM against FIPV 3CLpro in the FRET assay (Perera et al., 2018) and an EC_50_ value of 0.02~0.04 μM against the replication of FIPV in cell culture (Kim et al., 2016; Kim et al., 2015). Considering the *in vivo* efficacy of GC376 in cats with FIP (Kim et al., 2016; Pedersen et al., 2018), all tested compounds, especially GC543 and GC583, seem to have a good potential for further development as a drug for these viral infections. GC583 has been previously shown to have *in vivo* efficacy by significantly decreasing murine norovirus replication in the intestinal tract in mice [compound 16 in (Galasiti Kankanamalage et al., 2015)].

Notably, despite the sequence homology differences in the 3CLpros of RHDV and EBHSV, the 3D homology modelling showed that the catalytic site topology of these lagovirus 3CLpros appears to be highly conserved, which may explain the comparable potency of compounds against RHDV1 and 2 and EBHSV 3CLpros. The preference of Cha over Leu at the P2 position is also shared among many caliciviruses, including lagoviruses, human norovirus and feline calicivirus. The crystal structures of human norovirus 3CLpro bound to GC376 or GC583 have previously revealed that Cha at the P2 position is filling the hydrophobic S2 subsite of 3CLpro more tightly than Leu (Chang et al., 2019), which is also predicted in the homology model of 3CLpro of RHDV2, highlighting the similar structural requirements for 3CLpro inhibition of caliciviruses.

In summary, we assessed the potency of select protease inhibitors against RHDV1 and 2 and EBHSV using the *in vitro* assays and conducted structure-activity relationship and 3D homology modelling studies. Using the FRET and the cell-based reporter assays, we identified compounds that display potent inhibitory activities against all three lagovirus 3CLpros. To our knowledge, this is the first report on antiviral compounds that target 3CLpro for development of effective inhibitors broadly acting against pathogenic lagovirus.

## Acknowledgements

The authors would like to thank David George for technical assistance.

## Funding

This work was generously supported by USDA-NIFA AFRI (2019-67015-29864) and National Institutes of Health (R01 AI161085).

## Notes

### Competing Interest Statement

Kansas State University and Wichita State University have jointly filed a patent application that covers compounds in the manuscript with KOC, YK, and WCG as coinventors.

